# Combinatorial Detection of Conserved Alteration Patterns for Identifying Cancer Subnetworks

**DOI:** 10.1101/369850

**Authors:** Ermin Hodzic, Raunak Shrestha, Kaiyuan Zhu, Kuoyuan Cheng, Colin C. Collins, S. Cenk Sahinalp

**Affiliations:** Laboratory for Advanced Genome Analysis, Vancouver Prostate Centre, Vancouver, BC, Canada; School of Computing Science, Simon Fraser University, Burnaby, BC, Canada; Department of Urologic Sciences, University of British Columbia, Vancouver, BC, Canada; Department of Computer Science, Indiana University, Bloomington, IN, USA; Center for Bioinformatics and Computational Biology, University of Maryland, College Park, MD, USA

**Keywords:** conserved subnetwork, alterations, cancer, combinatorial optimization

## Abstract

**Background:** Advances in large scale tumor sequencing have lead to an understanding that there are combinations of genomic and transcriptomic alterations specific to tumor types, shared across many patients. Unfortunately, computational identification of functionally meaningful shared alteration patterns, impacting gene/protein interaction subnetworks, has proven to be challenging.

**Findings:** We introduce a novel combinatorial method, cd-CAP, for simultaneous detection of connected subnetworks of an interaction network where genes exhibit conserved alteration patterns across tumor samples. Our method differentiates distinct alteration types associated with each gene (rather than relying on binary information of a gene being altered or not), and simultaneously detects multiple alteration profile conserved subnetworks.

**Conclusions:** In a number of The Cancer Genome Atlas (TCGA) data sets, cd-CAP identified large biologically significant subnetworks with conserved alteration patterns, shared across many tumor samples.

## Introduction

Recent large scale tumor sequencing projects such as PCAWG (Pan Cancer Analysis of Whole Genomes) have revealed multitude of somatic genomic, transcriptomic, proteomic and epigenomic alterations across cancer types. However, only a select few of these alterations provide proliferative advantage to the tumor and hence are called “driver” alterations [1]. Distinguishing driver alterations from functionally inconsequential random “passenger” alterations is critical for therapeutic development and cancer treatment.

Cancers are often driven by alterations to multiple genes [2,3]. Whereas genomic alterations are likely consequences of endogenous or exogenous mutagen exposures [4], their evolutionary selection depends on the functional role of the affected genes [1] and their synergistic combinations. For example, *TMPRSS_2_-ERG* gene fusion is an early driver event in almost half of prostate cancer cases, and it often co-exists with copy-number loss of *PTEN* and *NKX_3-1_* [5, 6, 7]. Another example is the concomitant deletion of four cancer genes - *BAP_1_, SETD_2_, PBRM_1_*, and *SMARCC_1_* in chromosome locus 3p21, identified as a driver event in clear cell renal cell carcinoma (ccRCC) [8], uveal melanoma [9], and mesotheliomas [10]. These genes are involved in chromatin remodeling process, and their loss further impairs DNA damage repair pathway in tumors [9].

Alterations in two or more genes might be evolutionary co-selected because alteration in one gene might enhance the deleterious effect of the others [11]. Such co-selected genes are often active in a functionally significant subnetwork (i.e. module or pathway) within the human gene/protein interaction network and aberrations in such subnetworks are common to particular cancer types as demonstrated by recent sequencing efforts (e.g. PCAWG) [12]. For instance, *TMPRSS_2_* interacts with *ERG* and *PTEN* (see the example above) in STRING v.10 protein-protein interaction network; in fact all three genes co-operate to modulate NOTCH signaling pathway in *TMPRSS_2_-ERG* positive prostate cancer progression [7]. As a result, it is desirable to identify subsets of functionally interacting genes which are commonly (genomically or transcriptomically) altered in specific tumor types.

Recently, a number of computational methods have been developed to identify recurrent genomic (as well as transcriptomic) alteration patterns across tumor samples. Some of these methods have been designed to identify multiple gene alterations simultaneously based on their co-occurrence or mutual exclusivity relationships in a tumor cohort, either with [13] or without [14, 15] reference to a molecular interaction network. Other methods have been developed to identify subnetworks within a molecular interaction network with specific characteristics, e.g. the subnetwork of a fixed size with the highest total “weight” [16,17] or the subnetwork seeded by a particular node that can be derived through a diffusion process [18, 19]; naturally these methods do not capture recurrent alteration patterns across a cohort. A direction particularly relevant to our paper is motivated by a number of related works [18, 20, 21, 22], and explored by Bomersbach *et al.* [23], which suggests to find a subnetwork of a given size *k* with the goal of maximizing *h*, the number of samples for which at least one gene of the subnetwork is in an altered state. (A similar formulation where the goal is to maximize a weighted difference of *h* and *k*, for varying size *k*, can be found in [24].) Although the above combinatorial problems are typically NP-hard, they became manageable through the use of state of the art integer linear programming (ILP) solvers or greedy heuristics, or by the use of complex preprocessing procedures which significantly reduce the problem size.

Complementary to the ideas proposed above, there are also several approaches to identify mutually exclusive (rather than jointly altered) sets of genes and pathways [25, 26, 27]. These approaches utilize the mutational heterogeneity prevalent in cancer genomes, and are driven by the observation that mutations acting on same pathway are often mutually exclusive across tumor samples. Although, from a methodological point of view, these approaches are very interesting, they are not trivially extendable to the problem of identifying co-occurring alteration patterns (involving more than two genes) conserved across many samples.

### Our Contributions

In this paper we present a novel computational method, cd-CAP (combinatorial detection of Conserved Alteration Patterns), for detection of subnetworks of an interaction network, each with an alteration pattern conserved across a large subset of a tumor sample cohort. The framework of cd-CAP allows each gene to be labeled (or “colored”) with one or more distinct alteration types (e.g. somatic mutation, copy number alteration, or aberrant expression) with the goal of identifying one or more subnetworks, each with a specific alteration (labeling) pattern, that is shared across many samples (Figure 1). As such, cd-CAP solves a novel problem that has not been tackled in the literature. In fact, the very notion of *conserved subetworks* used by cd-CAP is novel: in [23,24] the subnetworks of interest are composed of nodes such that in each patient at least one is altered (one way or another). In contrast, cd-CAP insists that each node is altered in each patient, and each node preserves its alteration type in each patient. Additionally, unlike [24] which employ heuristics to solve a highly restrictive problem and thus cannot guarantee optimality, cd-CAP uses a very efficient exhaustive search method (a variant of the a-priori algorithm, originally designed for association rule mining [28]) to quickly solve a very general problem optimally.

**Figure 1.**
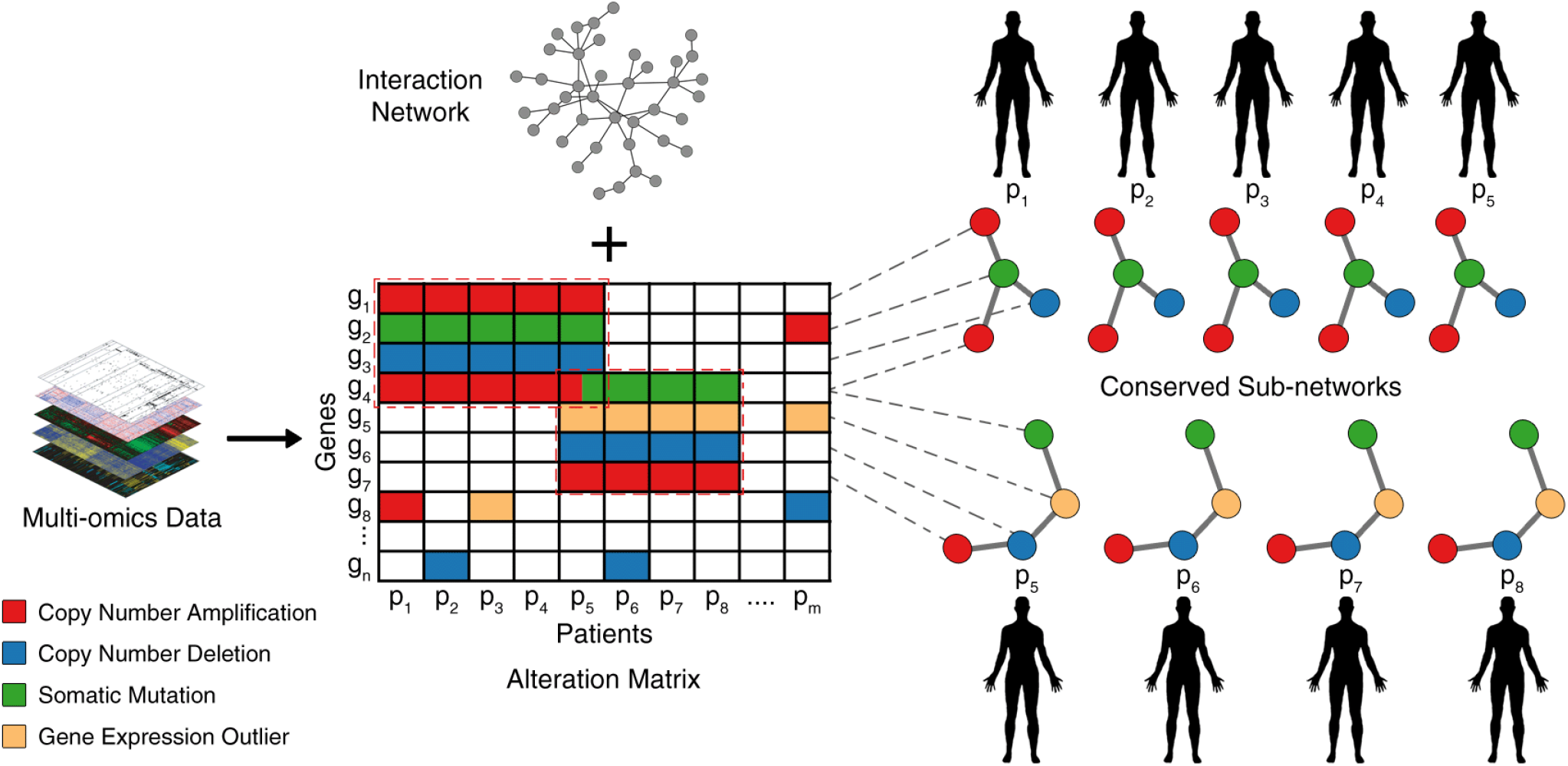
Schematic Overview of our framework. Multi-omics alteration profiles of a cohort of tumor samples are identified using appropriate bioinformatics tools. The alteration information is combined with gene-level information in the form of a sample-gene alteration matrix. Each alteration type is assigned a distinct color. Using a (signaling) interaction network, cd-CAP identifies subnetworks with conserved alteration patterns.

cd-CAP offers two basic modes: the “single-subnetwork” mode identifies the largest subnetwork altered the same way in at least *t* samples by solving the *maximum conserved subnetwork identification* problem optimally; the “multi-subnetwork” mode identifies *l* subnetworks of size (at most) *k* (*k* and *l* are user defined parameters) that collectively cover the maximum number of nodes in all samples by solving the *maximum conserved subnetwork cover* problem via ILP. In both modes, cd-CAP runs in two steps. The first step computes a set of all “candidate” subnetworks (each with a distinct alteration pattern) with at most *k* nodes, and which are shared by at least *t* samples. However, the two modes differ in the second step: the first returns a single largest subnetwork, and the second returns *l* subnetworks collectively covering the maximum number of nodes from the set of candidate subnetworks.

Additionally cd-CAP provides the user the ability to add or relax some constraints on the subnetworks it identifies. Specifically, the user can ask cd-CAP to (i) return “colorful” subnetworks (i.e. subnetworks of nodes with at least two distinct colors), or (ii) allow up to a *δ* fraction of nodes in the subnetwork to have no alteration (as a result, not colored) in some of the samples that share the subnetwork.

We have applied cd-CAP-with both single and multi-subnetwork mode, with the basic setting (which only requires that each node has the same alteration type across the samples), as well as each of the possible additional options above, i.e., (i), (ii) - to The Cancer Genome Atlas (TCGA) breast adenocarcinoma (BRCA), colorectal adenocarcinoma (COAD), and glioblastoma multiforme (GBM) datasets. On these datasets, which collectively include > 1000 tumor samples, cd-CAP identified several connected subnetworks of interest, each exhibiting specific gene alteration pattern across a large subset of samples.

In particular, cd-CAP results with the basic setting demonstrated that many of the largest highly conserved subnetworks within a tumor type solely consist of genes that have been subject to copy number gain, typically located on the same chromosomal arm and thus likely a result of a single, large scale amplification. One of these subnetworks cd-CAP observed (in about one third of the COAD samples [29]) include 9 genes in chromosomal arm 20q, which corresponds to a known amplification recurrent in colorectal tumors. Another copy-number gain subnetwork cd-CAP observed in breast cancer samples corresponds to a recurrent large scale amplification in chromosome 1 [30]. It is interesting to note that cd-CAP was able to re-discover these events without specific training.

Several additional subnetworks identified by cd-CAP solely consist of genes that are aberrantly expressed. Further analysis with option (ii) in the multi-subnetwork mode of cd-CAP revealed subnetworks that capture signaling pathways and processes critical for oncogenesis in a large fraction of tumors. We have also observed that the subnetworks identified through all different options of cd-CAP are associated with patients’ survival outcome and hence are clinically important.

In order to assess the statistical significance of subnetworks discovered by cd-CAP - in the single-subnetwork mode, we introduce for the first time a model in which likely interdependent events, in particular amplification or deletion of all genes in a single chromosome arm, are considered as a single event. Conventional models of gene amplification either consider each gene amplification independently [31] (this is the model we implicitly assume in our combinatorial optimization formulations, giving a lower bound on the true p-value), or assumes each amplification can involve more than one gene (forming a subsequent sequence of genes) but with the added assumption that the original gene structure is not altered and the duplications occur in some orthogonal “dimension” [32, 33, 34]. Both models have their assumptions that do not hold in reality but are motivated by computational constraints: inferring evolutionary history of a genome with arbitrary duplications (that convert one string to another, longer string, by copying arbitrary substrings to arbitrary destinations) is an NP-hard problem (and is difficult to solve even approximately) [35, 36]. By considering all copy number gain or loss events in the same chromosomal arm as a single event, we are, for the first time, able to compute an estimate that provides an empirical upper bound to the statistical signifl-cance (p-value) of the subnetworks discovered. (Note that this is not a true upper bound since a duplication event may involve both arms of a chromosome - but that would be very very rare.) Through this upper bound, together with the lower bound above, we can sandwich the true p-value and thus the significance of our discovery.

The “Combinatorial Optimization Formulation” section below describes the combinatorial optimization formulations used by cd-CAP to solve the problem of detecting conserved alteration patterns and all its above mentioned variants. The “Algorithmic Details” section describes implementation details for the two main steps of cd-CAP’s solution. The “Additional Constraints and Parameter Options” subsection describes the implementation details for the two variants on the constraints imposed by cd-CAP.

## Methods

### Combinatorial Optimization Formulation

Consider an undirected and node-colored graph *G* = (*V, E*), representing the human gene or protein interaction network, with *n* nodes where *v_j_* ∈ *V* represent genes and *e* = (*v*_*h*_, *v*_*j*_) ∈ *E* represent interactions among the genes/proteins. A given sample/patient *P_i_* (among *m* samples in a cohort) has a specific coloring of *G*, namely *G_i_* = (*V, E, C_i_*), where each node *v_i,j_* (corresponding to node *v_j_* ∈ *V*) is colored with one or more possible colors to form the set *C_i,j_* (i.e. *C_i_* maps *v_i,j_* to a possibly empty subset of colors *C_i,j_*). Each color represents a distinct type of alteration harbored by a gene/protein: specific alteration types we consider are somatic mutation (single nucleotide alteration or short indel), copy number gain, copy number loss or sig-niflcant alteration in expression (this set of alterations can be trivially expanded to include genic structural alteration - micro-inversion or duplication, gene fusion, alternative splicing, methylation alteration, non-coding sequence alteration) observed in a gene or its protein product. Note that *C_i,j_* = ø implies none of the alteration types we consider are observed at *v_i,j_*. Also note that given a node *v_j_*, its occurrences *v_i,j_* and *v*_*i’,j*_, in respective samples *P_i_* and *P’_i_*, have at least one matching color if *C_i, j_* ∩ *C*_*i’,j*_ ≠ ø.

The main goal of cd-CAP is to identify conserved patterns of (i.e. identically colored) connected subnetworks across a subset of colored (sample) networks *G*_*i*_. Consider a connected subnetwork *T* = (*V*_*T*_, *E*_*T*_) of the interaction network *G*, where each node *v_j_* ∈ *V_T_* is assigned a single color *c_j_*. Such a colored subnetwork is said to be shared by a collection of patient networks {*G*_*i*_: *i* ∈ *I*} if the color *c_j_* assigned to each vertex *v_j_* is in the color set *C_i,j_* of each *v_i,j_*(*i* ∈ *I*), i.e. *c_j_* ∈ ∩*_i∈I_C_i,j_* for each *v_j_* ∈ *V_T_*. Note that *v_i,j_* is said to be covered by a colored subnetwork if that colored subnetwork is shared by *G_i_* (Figure 1). Intuitively, a colored subnetwork represents a conserved pattern or a network motif.

In the single-subnetwork mode, cd-CAP solves the **Maximum Conserved Subnetwork Identification problem (MCSI)**, a specific combinatorial problem to identify conserved patterns of subnetworks. **MCSI** asks to find the largest connected colored subnetwork *S* of the interaction network *G*, that occurs in exactly *t* (a user specified number) samples 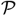, such that each node in *S* has the same color in each sample 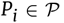. Note that this formulation is orthogonal to that used in [23] and [24], where the goal is to maximize the number of samples that share a fixed size subnetwork. Unlike these formulations, **MCSI** admits a generalization of the a-priori algorithm, which we use to solve it efficiently. Note that our formulation considers distinct types of mutations (as colors) in the conserved alteration patterns, another key improvement to alternative formulations used in the literature [23, 24].

In the multi-subnetwork mode, on the other hand, cd-CAP aims to simultaneously identify multiple conserved subnetworks that are altered in a large number of samples. In particular, it may aim to cover all nodes *v*_*i,j*_, in all *m* input sample networks *G*_*i*_, with the smallest number of subnetworks *T* = (*V*_*T*_*, *E*_*T*_*) shared by at least one sample network. We refer to this combinatorial optimization problem as **Minimum Subgraph Cover Problem for (Node) Colored Interaction Networks (MSC-NCI)**. As will be shown below, cd-CAP solves a slightly more constrained variant of this problem in the multi-subnetwork mode.

The MSC-NCI problem, as described above, is parameter-free. However, in a realistic multi-omics cancer dataset, the number of genes far exceeds the number of samples represented. Under such conditions, the solution to the MSC-NCI problem will primarily include subnetworks that are large connected components that are shared by only one sample network. To account for this situation, we introduce the following parameters/constraints akin to those for the MCSI formulation: (1) we require that the nodes in each subnetwork have their assigned color shared by at least *t* samples, (in the remainder of the discussion, *t* is referred to as *depth* of a subnetwork); and (2) we require that each subnetwork returned contains at most *k* nodes. Note that this variant of the problem is infeasible for certain cohorts (consider a particular node which has a unique color for a particular sample; clearly requirement (1) can not be satisfied if *t* >1). Even if there is a feasible solution, the requirement that each subnetwork in 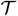 is of size at most *k* makes the problem NP-hard (the reduction is from the problem of determining whether *G* can be exactly partitioned into connected subnetworks, each with *k* nodes [37]). As a result (3) we introduce one additional parameter, *l*, the maximum number of subnetworks (each of size at most k, and which are color-conserved in at least *t* samples) with the objective of covering the maximum number of nodes across all samples. We call the problem of identifying at most *l* subnetworks of size at most *k*, whose colors are conserved across at least *t* samples, so as to maximize the total number of nodes in all these samples covered by these subnetworks, as the **Maximum Conserved Subnetwork Coverage problem (MCSC).**

### Algorithmic Details

In this section we describe the detailed algorithmic framework of cd-CAP, which consists of two steps for both its single and multi-subnetwork modes. The key insight as the basis of our algorithm is that in all instances of interest, only a limited number of genes are colored in comparison to the total number of nodes *nm*. This enables us to apply an exhaustive search method that is designed for association rule mining [28] to build a list of all “candidate subnetworks” exactly and efficiently (e.g. in comparison to the ILP or heuristic solutions in [23, 24]). Note that our exhaustive search method is an extension of the a-priori algorithm with the difference that we require the candidate subnetworks to maintain connectivity as they grow. As a result, we first compute the candidate subnetworks (each with a distinct alteration pattern) with at most *k* nodes, and which are shared by at least *t* samples in both modes. In the next step, in the single-subnetwork mode, cd-CAP simply returns the largest subnetwork among the candidate subnetworks, while in the multi-subnetwork mode it solves the maximum coverage problem (MCSC) on the set of candidate subnetworks via the ILP formulation below.

#### First step of cd-CAP: Generating candidate subnetworks

We generate the complete list of candidate subnetworks with minimum depth *t* by the use of *anti-monotone* property [38]: if any subnetwork *S* has depth < *t*, then the depth of all of its supergraphs *S’* ⊃ *S* must be < *t*. This makes it possible to grow the set *S* of valid subnetworks comprehensively but without repetition (described as “optimal order of enumeration” in [39]) through the following breadth-first network growth strategy.

(1) For every colored node *v_i,j_* and each of its colors *c_ℓ_*, we create a candidate subnetwork of size 1 (i.e. with single node) containing the node with color *c_ℓ_*. All samples in which the node is colored *c_ℓ_* trivially share this subnetwork.
(2) We inductively consider all candidate subnetworks of size *s* with the goal of growing them to subnetworks of size *s* + 1 as follows. For a given subnetwork *T* of size *s*, consider each neighboring node *u*. For each possible color 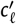. of *u*, we create a new candidate subnetwork of size *s* + 1 by extending *T* with *u* - with color 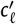. We maintain this subnetwork for the next inductive step only if the number of samples sharing this new subnetwork is at least *t*; otherwise, we discard it.

Once the procedure terminates, the single-subnetwork mode simply returns all subnetworks constructed in the final iteration (of size *s*). The multi-subnetwork mode requires additional processing as will be described below. Note, however, that during the extension of *T* above, if the new node *u* does not reduce the number of samples sharing it, *T* becomes redundant and is not considered in the ILP formulation

#### Second step of cd-CAP: Solving MCSC for multi-subnetwork mode

Given the universe *u* = {*v_i,j_* | *C_i,j_* ≠ ø, *i* = 1, · ·, *m*; *j* = 1, …, *n*}, containing all the colored nodes in all the sample networks, and the collection of all subnetworks *S* = {*T_i_* | *T_i_* is shared by ≥ *t* samples & contains ≤ *k* nodes}, our goal is to identify up to *l* subnetworks from the set *S* which collectively contain the maximum possible number of elements of the universe *u*.

After the list of all candidate subnetworks *S* is constructed (as described in the previous subsection), we represent the MCSC problem with the integer linear program below and solve it using IBM ILOG CPLEX or Gurobi. A binary variable *C*[*i,j*] corresponds to whether colored node *v_i,j_* was covered by at least one chosen subnetwork, and binary variable *X*[*i*] corresponds to whether colored candidate subnetwork *T_i_* was one of the chosen. Similarly *S_i,j_* represents the set of all subnetworks of *S* which contain node *v_i,j_* properly colored in them.

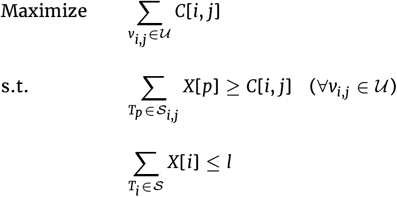

### Additional Constraints and Parameter Options

In addition to the exactly-conserved colored subnetworks obtained through the general MCSI or MCSC formulation as described above, cd-CAP offers the user to add or relax constraints through new parameters, in both single and multisubnetwork mode.

i. *“Colorful” Conserved Subnetworks*. In some of the datasets that we analyzed, certain variant types (i.e. colors) were dominant in the input to an extent that all subnetworks identified by our method had all nodes colored identically. By insisting that the identified subnetworks are *colorful*, it is possible to, e.g., capture conserved genomic alterations and their impact on their interaction partners (form of expression alterations). For this purpose we introduce the notion of a *colorful subnetwork, T*, as a subnetwork that has at least two distinct colors represented in the coloring of its nodes, i.e. 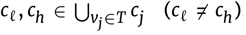. In order to identify colorful subnetworks instead of arbitrary subnetworks, we update the first step of cd-CAP so that it specifl-cally keeps track of colorful subnetworks (rather than all subnetworks) in each iteration; this is because any colorful network must contain a connected colorful subnetwork.
ii. *Subnetworks Conserved within error rate *δ**. In order to reduce the sensitivity of cd-CAP to noise (that emerges during the assignment of variant types to genes - due to limited precision of sequence or statistical analysis methods) in the input data, we provide the user the option to allow *errors* in identifying conserved subnetworks. For that, cd-CAP provides the user the option to specify an error rate *δ* that represents the fraction of nodes in a subnetwork *T* that can have no assigned color in any sample that shares *T*. We implemented this by updating the first step of cd-CAP so that it expands the set of samples that share each candidate subnetwork *T* to every other sample where *T* occurs with ≤ *δ*|*T*| color omissions.

### Assessing the Statistical and Biological Significance of the Networks Identified by cd-CAP

#### Statistical significance of subnetworks identified by cd-CAP

It is possible to assess the statistical significance of the subnetworks identified by cd-CAP by applying the conventional permutation test [13, 23, 27] on the color assignments of nodes - under the assumption that each gene is altered independently: let *C_i,j_* represent the set of colors assigned to a node *v_i,j_* and let *C_i_* = {(*v*_*i,j*_,*C*_*i,j*_)}, represent the entire set of color assignments to nodes *v_i,j_* in network *G_i_*. We can obtain a random permutation of the color assignment 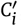, by independently shuffiing each color *c* ∈ ∪*_j_C_i,j_* across the nodes of *G_i_*, which results in an assignment of a new color set 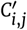 to each node *v*_*i,j*_, under the constraint that the total number of nodes with each color *c* is preserved. For a subnetwork *T* = (*V*_*T*_, *E*_*T*_) of size *k* covering *t* samples returned by cd-CAP in the single-subnetwork mode, we can carry out a permutation test as follows. First we generate a permuted color assignment (as described above) for each sample. Then we run cd-CAP in the single-subnetwork mode (possibly with the option (i) or (ii) as described in the previous section) and identify the largest subnetwork which covers at least *t* samples. We repeat this sufficiently many (by default 1000) times to compute *P*_1,*T*_, the number of times we end up with a subnetwork of size at least *k* in *t* or more samples, normalized by the number of attempts. We can use *P*_1,*T*_ as an empirical p-value for subnetwork *T* of size *k*.

*P*_1,*T*_ forms an empirical lower bound for the p-value of *T* rather than an accurate estimate since it ignores the interdependencies among gene alteration events (i.e. node colors). In particular, whole chromosome or chromosome arm level copy number amplifications/deletions are commonly observed in cancer - such events must be reflected in the permutation test we employ. To address this issue, we apply the following procedure to compute *P*_2_,*_T_* as an empirical upper-bound for the p-value of *T*, under the assumption that copy number alterations take place in whole chromosome arms. For a given color *E*, corresponding to either copy number gain or loss events, let *N_i,E_* denote the number of nodes with color *E* in *G_i_*. For each chromosomal arm *A*, consider the set of nodes *v_i,A_* that have been assigned at least one color in *G_i_*. Now we can reassign colors to vertices such that (1) colors *E* corresponding to copy number gain or loss are assigned to all genes in a chromosome arm simultaneously; specifically the set of nodes *V*_*i*_ in a chromosome arm *A* are all assigned the same color *E* independently with probability 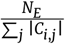 (which guarantees that the expected number of nodes with color *E* in *G_i_* is preserved); (2) the remaining colors (not related to copy number gain or loss) are assigned randomly to those nodes without a color assignment thus far (as described in the computation for *P*_1,*T*_). This process provides a new randomly permuted color assignment 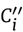 which we use to obtain an empirical upper bound on the p-value of a subnetwork *T* discovered by cd-CAP. For that perform this process simultaneously in all *G_i_* and check whether the largest subnetwork shared by at least *t* samples exceeds the size of a subnetwork *T* (identified on the input dataset by cd-CAP). We repeat this process sufficiently many times and record the number of times the largest subnetwork obtained indeed exceeds the size of *T*; that value normalized by the number of times the process is executed is the value *P*_2,*T*_, the empirical upper bound on the p-value of *T*. The true p-value of *T* must be in the range [*P*_1,*T*_, *P*_2,*T*_] (provided that chromosome arms form the largest units of alteration).

#### Pathway enrichment analysis

We tested the set of genes in the subnetworks obtained by cd-CAP for enrichment against gene sets corresponding to pathways present in the Molecular Signature Database (MSigDB) v6.0 [40]. A hypergeometric test based gene set enrichment analysis [40] was used for this purpose. A false discovery rate (FDR) ≤ 0.01 was used as a threshold for identifying significantly enriched pathways.

#### Association between cd-CAP identified sub-networks and patients’ survival outcome

In order to assess the association between each cd-CAP identified subnetwork *T* with patients’ survival outcome, we used a risk-score based on the (weighted) aggregate expression of all genes in the subnetwork *T*. The risk-score (*S*) of a patient is defined as the sum of the normalized gene-expression values in the subnetwork, each weighted by the estimated univariate Cox proportional-hazard regression coefficient [41], i.e., 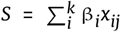 Here *i* and *j* represent a gene and a patient respectively, ß_i_ is the coefficient of Cox regression for gene *i, x_ij_* is the normalized gene-expression of gene *i* in patient *j*, and *k* is the number of genes in the subnetwork. The normalized gene-expression values were fitted against overall survival time with living status as the censored event using univariate Cox proportional-hazard regression (exact method). Based on the risk-score values, patients were stratified into two groups: low-risk group (patients with *S* < mean of *S*), and high-risk group (patients with *S* ≥ mean of *S*). Note that only those patients that are covered by the subnetwork are considered for the analysis above. In fact, with respect to survival outcomes, the set of patients covered by a subnetwork identified by cd-CAP would not necessarily differ from those that are not, since the latter set is likely to be highly heterogeneous with respect to cancer subtypes.

## Results

### Datasets and data processing

#### TCGA tumor variant data

We obtained somatic mutation, copy number aberration and RNA-seq based gene-expression data from three distinct cancer types - glioblastoma multiforme (GBM) [42], breast adenocarcinoma (BRCA) [43], and colon adenocarcinoma (COAD) [29] from The Cancer Genome Atlas (TCGA) datasets (detailed information can be found in Supplementary Section 1). In addition, we distinguish four commonly observed molecular subtypes (i.e. Luminal A, Luminal B, Triple-negative/basal-like and HER2-enriched) from the BRCA cohort. For each sample, we obtained the list of genes which harbor somatic mutations, copy number aberrations, or are expression outliers as per below.

#### Somatic Mutations

All non-silent variant calls that were identified by at least one tool among MUSE, MuTect2, Somatic-Sniper and VarScan2 were considered.

#### Copy Number Aberrations

CNA segmented data from NCI-GDC were further processed using Nexus Copy Number Discovery Edition Version 9.0 (BioDiscovery, Inc., El Segundo, CA) to identify aberrant regions in the genome. We restricted our analysis to the most confident CNA calls selecting only those genes with high copy gain or homozygous copy loss.

#### Expression outliers

We used HTSeq-FPKM-UQ normalized RNA-seq expression data to which we applied the generalized extreme studentized deviate (GESD) test [44]. In particular, we used GESD test to compare the transcriptome profile of each tumor sample (one at a time) with that from a number of available normal samples. For each gene, if the tumor sample was identified as the most extremely deviated sample (using critical value α = 0.1), the corresponding gene was marked as an expression-outlier for that tumor sample. This procedure was repeated for every tumor sample. Finally, comparing the tumor expression profile of these outlier genes to the normal samples, their up or down regulation expression patterns were determined.

#### Interaction networks

We used the following human protein-interaction networks in the identification of the most significant subnetworks specific to the cancer types mentioned above. (1) STRING version 10 [45] protein-interaction network which contains high confidence functional protein-protein interactions (PPI). Self-loops and interactions with missing HGNC symbols were discarded and interaction scores were normalized (divided by 1000) to obtain a reliability score in the range [0, 1]. Only high confidence interactions with combined score of 0.9 or greater were selected. (2) STRING network with only experimentally verified edges. (3) Human Protein Reference Database (HPRD) version 9 [46]. (4) REACTOME version 2015 [47].

### Maximal Colored Subnetworks Across Cancer Types

We used cd-CAP to solve the maximum conserved subnetwork identification (MCSI) problem exactly on each of the protein-interaction networks we considered on all cancer types - for every feasible value of network depth. As can be easily observed, the depth and the size of the identified subnetwork are inversely related. We say that a network depth value is feasible if (i) the depth is at least 10% of the cohort size, (ii) the maximum network size for that depth is at least 3, (iii) the number of “candidate” subnetworks are at most 2 millions per iteration when running cd-CAP for that depth.

The number of maximal solutions of cd-CAP as a function of feasible network depth for each cancer type (COAD, GBM, BRCA Luminal A, and BRCA Luminal B) is shown in Figure 2A-D on STRING v10 PPI network with high confidence edges (see Supplementary Figure 2–5 for the results on alternative PPI networks). In general, for a fixed network size, the number of distinct networks of that size decreases as the network depth increases. One can observe that the end of “valleys” in the colored plots in Figure 2A-D correspond to the largest depth that can be obtained for a given subnetwork size.

**Figure 2.**
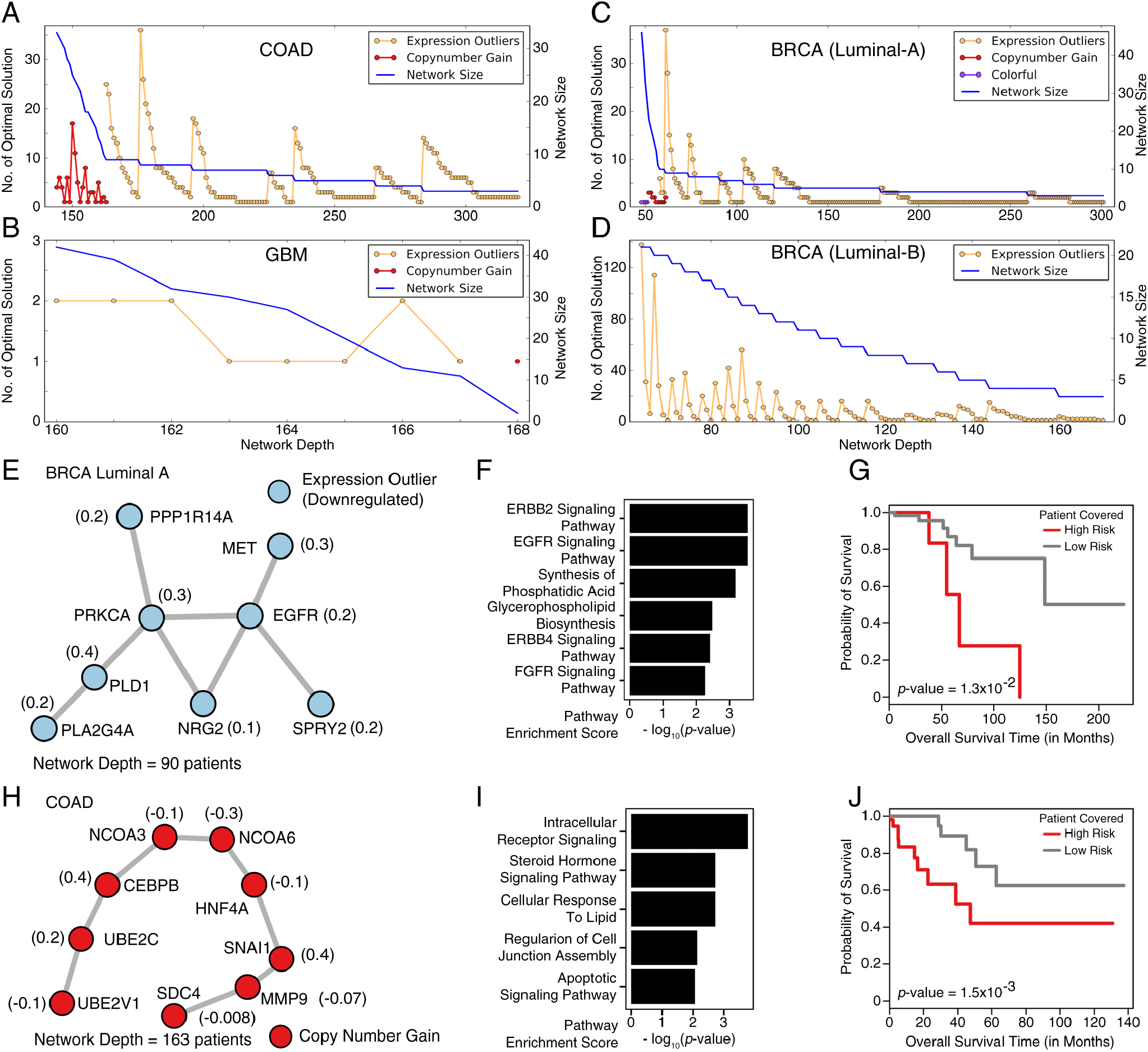
Conserved colored subnetworks. (A-D) Number of maximal solutions and the size of the conserved colored subnetwork obtained using the MCSI formulation, as a function of network depth *t*, in each of four cancer types analyzed, on STRING v10 (with high confidence edges) PPI network. The horizontal axis denotes the depth (number of patients) of the network. For the blue plot, the vertical axis denotes the maximum possible network size (in terms of the number of nodes) and thus it is strictly non-increasing by definition. For the plots with different colors, the vertical axis denotes the number of distinct networks with network size equal to that indicated by the blue plot. As can be seen, the red plots depict networks where all nodes have a copy number gain, the yellow plots depict networks where all nodes are expression outliers and purple plots depict colorful networks (with at least two distinct colors). A total of 41 subnetworks across all cancer types (10 COAD, 4 GBM, 11 Luminal A and 16 Luminal B) correspond to the end of “valleys” in the color plots - and were further analyzed. Two of the most interesting ones are provided here - both of which are uni-colored. The number in parenthesis next to each node represents the univariate Cox proportional-hazard regression coefficient estimated for each gene, used as its weight in the risk-score calculation to stratify patients into two distinct risk groups. (See Methods section for details). (E-G) One of the 11 maximal colored subnetworks identified in BRCA Luminal A dataset: it consists solely of downregulated expression outlier genes and has depth 90 (patients). (E) The colored subnetwork (with 8 nodes) topology. (F) Pathways dysregulated by alterations harboured by the genes in the subnetwork - these genes are involved in EGFR, ERBB2, and FGFR signaling pathways. (G) Kaplan-Meier plot showing the significant association of the subnetwork, with patients’ clinical outcome. Patients “covered” by the subnetwork were stratified into two groups, namely High Risk (8 patients) vs Low Risk (82 patients), based on their gene expression levels. (See Methods for details.) (H-J) One of the 10 maximal colored subnetworks identified in COAD dataset - it consists solely of copy number amplified genes and has a depth of 163 (patients). Genes in this subnetwork belong to the same chromosomal locus 20q13. (H) The colored subnetwork (with 9 nodes) topology. (I) Pathways dysregulated by the alterations harboured by the genes in the subnetwork - these genes are involved in signal transduction and apoptotic process. (J) Kaplan-Meier plot showing the significant association of the subnetwork with patients’ clinical outcome (73 High Risk vs 83 Low Risk patients).

In the remainder of the paper we focus only on the single colored subnetwork of each given size that has the maximum possible depth (corresponding to the end of “valleys” in the plots). (If for a given subnetwork size and the corresponding maximal depth, cd-CAP returns more than 1 subnetwork, they are ignored.)

Many of the subnetworks we focused on, especially those with large depth, only consisted of expression outlier genes (typically all upregulated or all downregulated) (Figure 2A-D) - across all four cancer types. In Luminal A data set for example, cd-CAP identified a subnetwork of eight downregulated genes with a network depth 90 (Figure 2E) - consisting of genes *EGFR, PRKCA, SPRY_2_*, and *NGR_2_*, known to be involved in EGFR/ERBB2/ERBB4 signaling pathways (Figure 2F). *EGFR* is an important driver gene involved in progression of breast tumors to advanced forms [48] and its altered expression is observed in a number of breast cancer cases [30]. The subnetwork also included MET, another well-known oncogene [49], and is enriched for members of the Ras signaling pathway, which is also known for its role in oncogenesis and mediating cancer phenotypes such as over-proliferation [50].

cd-CAP additionally identified some (uni-colored) copy-number gain networks, typically with lower depth: a prominent example is in the COAD dataset with depth 163 (out of 463 patients in the cohort). This network forms the core of larger (maximal) subnetworks cd-CAP identifies for lower depth values; it corresponds to a copy number gain of the chromosomal arm 20q - a well known copy number aberration pattern highly specific to colorectal adenocarcinoma tumors [29]. Another subnetwork cd-CAP identified in 15% of the 422 BRCA Luminal A samples corresponds to a copy number gain on chromosome 1 which is again a known aberration associated with breast cancer [30].

Note that cd-CAP also identified several multi-colored subnetworks. The benefits of cd-CAP’s ability to identify multicolored subnetworks is demonstrated in Supplementary Figure 1, which summarizes the results of a comparison between cd-CAP and a limited version of cd-CAP that does not differentiate mutation types. The figure shows that, especially in COAD and GBM, the survival outcomes of samples that include the cd-CAP identified subnetworks differ significantly from those sameples that do not include such subnetworks. In the BRCA data set, since all subnetworks of interest involve differentially expressed genes, the difference between survival outcomes is insignificant.

A complete list of subnetworks of focus (from STRING v10 with high confidence edges), across all cancer data sets, is provided in the Supplementary Table 2. For each of these subnetworks, and for each patient covered by a particular subnetwork, we calculated a risk-score defined as a linear combination of the normalized gene-expression values of the genes in the subnetwork weighted by their estimated univariate Cox proportional-hazard regression coefficients (see Methods section for details). Based on the risk-score values, the patients covered by the subnetwork were stratified into two risk groups (high risk and low risk group).

The expression outlier subnetwork we mentioned above for the Luminal A dataset was the most significant among all subnetworks identified in this dataset (Figure 2G). The patients in the high-risk group have poor overall survival outcome suggesting clinical importance of the identified subnetwork by cd-CAP.

Another copy-number gain subnetwork shared among 163 patients in the COAD dataset (Figure 2H) was comprised of genes from chromosome locus 20q13 - likely indicating a single chromosomal amplification event. Intriguingly, these genes form a linear structure on the protein interaction network. Among them is a group of functionally related genes consisting of transcription factors and their regulators (genes *CEBPB,NCOA’s, UBE_2_’s*), which are known to be involved in the intracellular receptor signaling pathway (Figure 2I). *CEBPB* and *UBE_2_’s* are also involved in the regulation of cell cycle [51]. At the other end of the linear subnetwork, there are *MMP_9_* and *SDC_4_*, established mediators of cancer invasion and apoptosis [52, 53]. We also confirmed that these genes are highly predictive of the patients’ survival outcome (Figure 2J). All these results seem to support that cd-CAP identified subnetworks are functionally important with potential clinical relevance.

### Maximal Colorful Subnetworks Across Cancer Types

We used cd-CAP to solve the maximum conserved colored subnetwork identification problem - with at least two distinct colors (see Section “Additional Constraints and Parameter Options” for details), in each of the four protein-interaction networks we considered and on each cancer type. Again, cd-CAP was run with every feasible value (as defined above) of network depth. The number of maximal solutions of cd-CAP as a function of network depth for each cancer type (COAD, GBM, BRCA Luminal A, and BRCA Luminal B) is shown in Figure 3A-D on STRING v10 PPI network with high confidence edges (see Supplementary Figure 2–4 for the results on alternative PPI networks). Note that we pay special attention to subnetworks with at least one sequence altered gene (i.e. a gene that is somatically mutated or copy number altered) since the sequence alteration(s) may explain expression-level changes in the remaining genes of the subnetwork (Figure 3E provides such an example).

**Figure 3.**
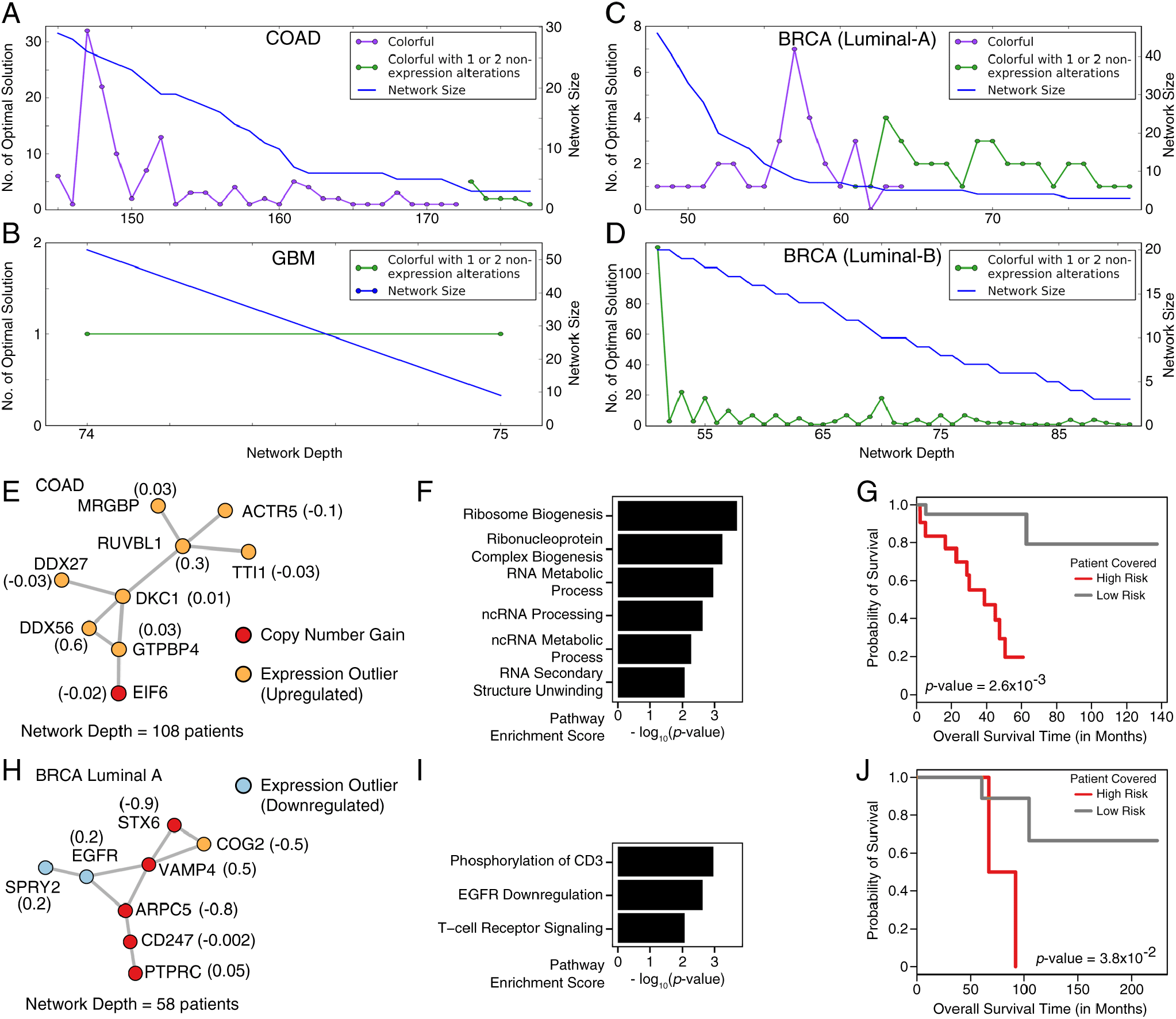
Colorful maximal subnetworks. (A-D) Number of maximal solutions and the size of the conserved colorful subnetwork obtained using the MCSI formulation, as a function of network depth t, in each of cancer types analyzed on the STRING v10 (high confidence edges) PPI network. The horizontal axis denotes the depth (number of patients) of the network. For the blue plot, the vertical axis denotes the maximum possible network size (in terms of the number of nodes) and thus it is strictly non-increasing by definition. For the plots with different colors, the vertical axis denotes the number of distinct networks with network size equal to that indicated by the blue plot. As can be seen, the purple plots depict colorful subnetworks and the green plots depict networks that include one to two nodes which are not expression outliers. A similar analysis was performed on the STRING v10 (experimentally validated edges), REACTOME and HPRD PPI networks. A total of 104 colorful subnetworks corresponding to the end of “valleys” of the plots were identified across the 4 cancer types in all the above PPI networks. Two of the most interesting ones are provided here. The number in parenthesis next to each node represents the univariate Cox proportional-hazard regression coefficient estimated for that gene, used as its weight in the risk-score calculation to stratify the patients into two distinct risk groups. (See Methods section for details). (E-G) One of the maximal colorful subnetworks identified in the COAD dataset, consisting of at most 2 non-expression outlier (for this case copynumber gain) genes, with depth 108 (patients). (E) The colored subnetwork (with 9 nodes) topology - obtained from STRING v10 (with experimentally validated edges) PPI network. (F) Pathways dysregulated by alterations harboured by the genes in the subnetwork - these genes are involved in Ribosome biogenesis and RNA processing. (G) Kaplan-Meier plot showing the significant association of the subnetwork, with patients’ clinical outcome (59 High Risk vs 47 Low Risk patients). (H-J) One of the maximal colorful subnetworks identified in the Luminal A dataset with no color restrictions, with depth of 58 (patients). (H) The colored subnetwork (with 8 nodes) topology - obtained in the REACTOME PPI network. (I) Pathways dysregulated by the alterations harboured by the genes in the subnetwork. (J) Kaplan-Meier plot showing the significant association of the subnetwork with patients’ clinical outcome (30 High Risk vs 30 Low Risk patients).

One such COAD subnetwork is composed of several overexpressed genes and one copy number gain gene - covering 108 patients (Figure 3E). This subnetwork is mainly enriched for genes involved in ribosome biogenesis (Figure 3G). Cancer has been long known to have an increased demand on ribosome biogenesis [54], and increased ribosome generation has been reported to contribute to cancer development [55]. The biological relevance of this subnetwork is also supported by survival analysis, which shows a strong differentiation between the high-risk and low-risk groups - see Figure 3F.

Another subnetwork we observed in 58 BRCA Luminal A samples consists of four copy number gain genes, an overexpressed gene, and two underexpressed genes, including *EGFR* (Figure 3H). All copy-number gain genes and the overexpressed gene are located in chromosome 1q, commonly reported in breast cancer [30]. The subnetwork involves an interesting combination of the down-regulation of the cancer gene *EGFR* and the amplification of a group of genes involved in T-cell receptor signaling (*PTPRC, CD_247_*, and *ARPC_5_;* see Figure 3I). Thus we may surmise that the covered population of patients potentially have relatively low cancer proliferation index with higher anti-tumor immune response, which can be highly relevant indicators with respect to clinical outcome. Indeed, this subnetwork is significantly associated with patients’ survival (Figure 3J).

### Multiple-Subnetwork Analysis Across Cancer Types

We next sought to detect up to 5 subnetworks per cancer type that collectively cover maximum possible number of colored nodes by solving the MCSC problem on STRING v10.5 network (with experimentally validated edges). The subnetwork extension error rate was set to 20%, and we restricted the search space to subnetworks which do not consist only of expression outlier nodes, in order to obtain what we believe to be more biologically interesting results. The network depth *t* was chosen for each dataset in a way that made it possible to construct all candidate subnetworks of maximum possible size while keeping the total number of candidate subnetworks below 2 × 10^6^, making the problem solvable in reasonable amount of time. We set *t* to 69 (15% of the patients), 62 (10% of the patients), and 110 (10% of the patients) respectively for COAD, GBM, and BRCA datasets. Supplementary Table 1 shows the size, per sample depth, and the coloring of the nodes in the resulting subnetworks.

We note that the subnetworks identified in the GBM dataset had the lowest depth (10-15% of the samples). COAD and BRCA datasets on the other hand have much larger depth (respectively 30-48% and 15-32% of the samples). Smaller subnetworks of the GBM dataset solely consist of copy number gain genes on chromosome 7q, a known amplification in GBM [56]. The two large subnetworks each contain a single gene with copy number gain (*SEC_61_G* and *EGFR*, respectively) accompanied by several of overexpressed genes. BRCA dataset exhibits a similar pattern: each of the four large subnetworks contain a single copy number gain gene from chromosome 8q, (*NSMCE_2_* in one and *MYC* in the remaining three subnetworks). Subnetworks detected in COAD dataset were much more colorful and recurrently conserved in a larger fraction of samples than those in the other datasets. All genes with copy number gain are located in chromosome 20q.

We identified a subnetwork with 15 nodes (11 genes with copy number gain, 1 overexpressed and 3 underexpressed genes) in 149 COAD patients (Figure 4A). All 11 copy number gain genes belong to chromosome 20q. *IL6R, PLCG_1_, PTPN_1_*, and *HCK* are involved in cytokine/interferon signaling to activate immune cells to counter proliferating tumor cells [57] (Figure 4B). *UBE_2_I, AURKA*, and *MAPRE_1_* are involved in cell cycle processes. This subnetwork was found to be associated with patients’ survival outcome (Figure 4C).

**Figure 4.**
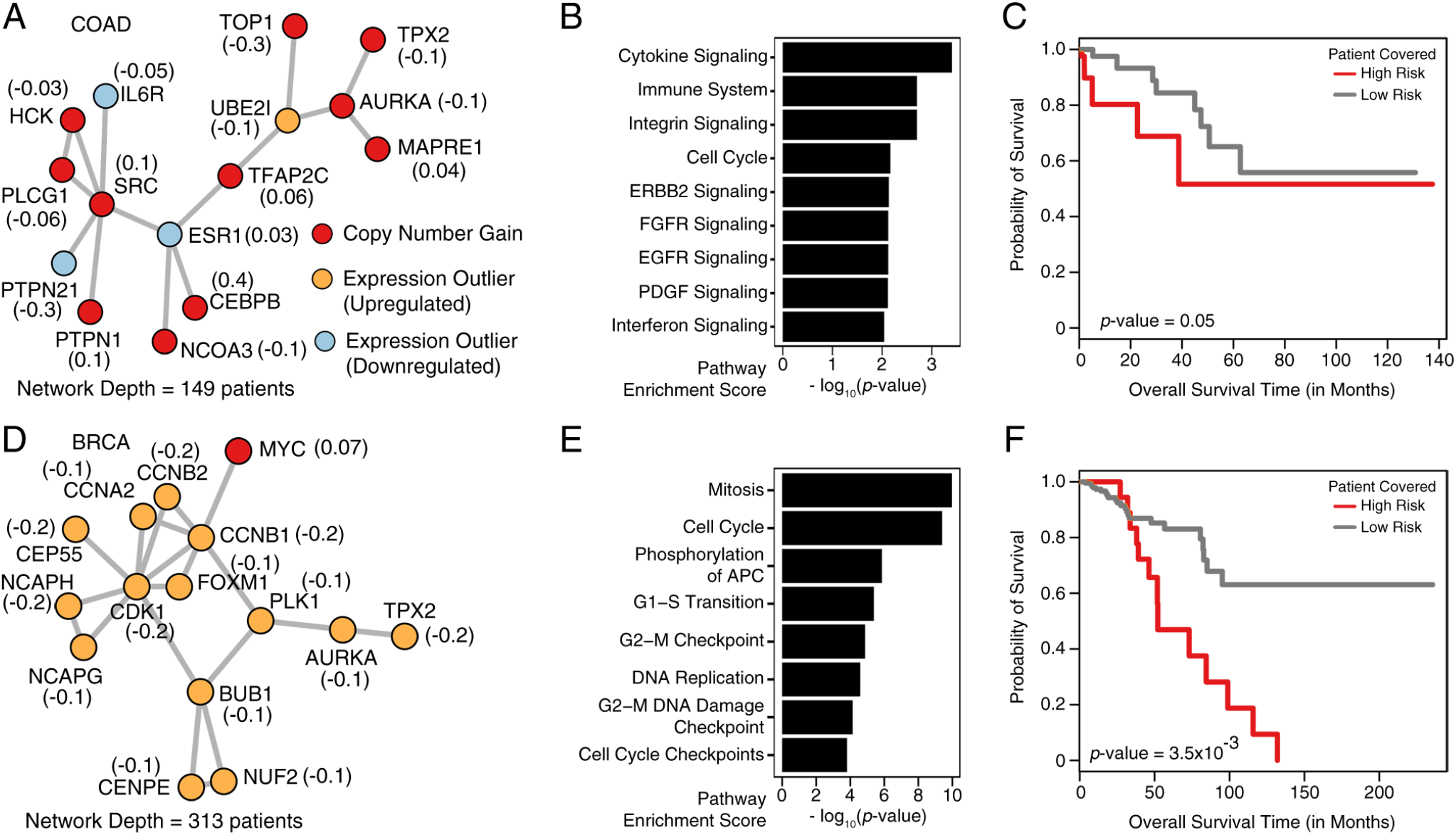
Multiple Subnetwork Analysis. Two of the largest subnetworks identified across the COAD, GBM and BRCA data sets (5 networks were identified per cancer type) through the MCSC formulation of cd-CAP on STRING v10.5 (with experimentally validated edges) PPI network. The number in parenthesis next to each node represents the univariate Cox proportional-hazard regression coefficient estimated for that gene, used as its weight in the risk-score calculation to stratify the patients into two distinct risk groups. (See Methods section for details). (A-C) The largest of the 5 COAD subnetworks with a network depth of 149 (patients). (A) The subnetwork topology (with 15 nodes). (B) Pathways dysregulated by alterations harboured by the genes in the subnetwork. (C) Kaplan-Meier plot showing the significant association of the subnetwork, with patients’ clinical outcome (69 High Risk vs 78 Low Risk patients). (D-F) The largest of the 5 BRCA subnetworks with a network depth of 313 (patients). (D) The subnetwork topology (with 15 nodes). (E) Pathways dysregulated by the alterations harboured by the genes in the subnetwork. (F) Kaplan-Meier plot showing the significant association of the subnetwork with patients’ clinical outcome (33 High Risk vs 278 Low Risk patients).

We identified another subnetwork with 15 nodes (14 overexpressed and 1 copy number gain genes) in 313 breast cancer patients (Figure 4D). Genes in this subnetwork are involved in cell cycle processes (Figure 4E). In particular the cell cycle checkpoint processes were dysregulated - which is known to drive tumor initiation processes [58]. The subnetwork was found to be associated with patients’ survival outcome (Figure 4F) demonstrating its clinical relevance.

### Empirical P-Value Estimates Confirm the Significance of cd-CAP Identified Networks

To evaluate the significance of cd-CAP’s findings, we performed the permutation test described earlier 1000 times on each cancer type for each possible setting of subnetwork constraints. Supplementary Table 2-3 and 0Figure 6 demonstrate the distribution of the empirical p-value upper bound estimates with STRING 10 (high confidence edges) PPI network, while the lower bound results look similar to what is presented in the figure and thus are omitted. In the permutation tests all cd-CAP identified subnetworks (without additional constraints) of size 2-5 were composed solely of expression altered genes; in contrast there are several larger CNV rich subnetworks observed in the TCGA COAD data set and others, further confirming the significance of our findings. Colorful subnetworks presented in Figure 3 are even less likely to occur at random (we therefore omit empirical p-value estimates for the networks in Figure 3).

## Discussion

In this paper we introduce a novel combinatorial framework and an associated tool named cd-CAP which can identify (one or more) subnetworks of an interaction network where genes exhibit conserved alteration patterns across many tumor samples. Compared with the state-of-the-art methods (e.g.[22, 24]), cd-CAP differentiates alteration types associated with each gene (rather than relying on binary information of a gene being altered or not), and simultaneously detects multiple *alteration type conserved* subnetworks.

cd-CAP provides the user with two major options. (a) In single-subnetwork mode, it computes the largest colored subnetwork that appears in at least *t* samples. This option exhibits significant speed advantage over available ILP-based approaches; its a-priori based algorithmic formulation allows flexible integration of special constraints (on maximal subnetworks) - not only simplifying complicated ILP constraints, but also further reducing the number of candidate subnetworks in iteration steps (a good example for this is the “colorful conserved subnetworks” as introduced in Section “Additional Constraints and Parameter Options”). However, the identified subnetworks are required to be *conserved*, i.e., each node only admits one alteration type among the samples sharing it (although we have relaxed constraints that allow each sample to have a few nodes without any alterations, i.e. colors). In the future, we may be able to extend the definition of a network to include nodes with color mismatches (for example, according to the definition in [21]) or [22] with a modification to cd-CAP’s candidate subnetwork generation algorithm. (b) In multi-subnetwork mode, it solves the *maximum conserved subnetwork cover* (MCSC) problem to cover the maximum number of nodes in all samples with at most *l* colored subnetworks (*l* is user defined) via ILP. In the future we aim to refine the MCSC formulation with reduced number of parameters and hope to develop exact or approximate solutions.

Subnetworks identified by cd-CAP in COAD, GBM and BRCA datasets from TCGA are typically enriched with genes harboring gene-expression alterations or copy-number gain. Notably, we observed that genes in subnetworks with copy-number amplification are universally located in the same chromosomal locus. Many of these genes have known interactions and are functionally similar, demonstrating the ability of cd-CAP in capturing functionally active subnetworks, conserved across a large number of tumor samples. These subnetworks seem to overlap with pathways critical for oncogenesis. In the datasets analyzed, we observed cell cycle, apoptosis, RNA processing, and immune system processes that are known to be dysregulated in a large fraction of tumors. cd-CAP also captured subnetworks relevant to EGFR/ERBB2 signaling pathways, which have distinct expression patterns in specific subtypes of breast cancer [30, 59]. Survival analysis of cd-CAP identified subnetworks also confirmed their substantial clinical relevance. In the future, it may be possible to use tissue-specific interaction data (such as [60] or [61]) to capture subnetworks with gene interactions that are more relevant to a specific cancer and tissue type.

## Availability of source code and requirements

- Project name: cd-CAP
- Project home page https://github.com/ehodzic/cd-CAP
- Operating system(s): Platform independent
- Programming language: C+ +
- Other requirements: make (version 3.81 or higher), g++ (GCC version 4.1.2 or higher), and IBM ILOG CPLEX Optimization Studio
- License: MIT License
- SciCrunch RRID: SCR_016843

## Declarations

**List of abbreviations**
BRCA: Breast Adenocarcinoma
ccRCC: Clear Cell Renal Cell Carcinoma
cd-CAP: combinatorial detection of Conserved Alteration Patterns
COAD: Colorectal Adenocarcinoma
CNA: Copy Number Aberrations
GBM: Glioblastoma Multiforme
GESD: Generalized Extreme Studentized Deviate
ILP: Integer Linear Programming
MCSI: Maximum Conserved Subnetwork Iden-tiflcation problem
MSC-NCI: Minimum Subgraph Cover Problem for (Node) Colored Interaction Networks
MCSC: Maximum Conserved Subnetwork Coverage problem
NCI-GDC: National Cancer Institute Genomic Data Commons
PCAWG: Pan Cancer Analysis of Whole Genomes
TCGA: The Cancer Genome Atlas

### Ethical Approval

Not applicable

### Consent for publication

Not applicable

### Author’s Contributions

S.C.S. conceived and directed the project. E.H., R.S., K.Z., and S.C.S. developed the algorithm. E.H. and K.Z. developed the cd-CAP software. E.H., R.S., K.Z., and K.C. performed the data analysis. R.S, K.C., and C.C.C. helped in biological interpretation of the results. All authors contributed to the preparation and revision of the manuscript.

## Acknowledgements

We thank Dr. Oktay Gunluk (IBM T. J. Watson Research Center) and the IBM Academic Initiative for providing us with free license to IBM ILOG CPLEX Optimization Studio (CPLEX). We thank all members of the Sahinalp and Collins laboratories for helpful suggestions.

