## Supplementary material for "Combinatorial Detection of Conserved Alteration Patterns for Identifying Cancer Subnetworks" (from another version of this preprint: version 2, 2018-12-13)

### cd-CAP: Supplementary Materials

#### 1 Data description

##### 1.1 Data Sets

The TCGA datasets were downloaded in February 2017 from the National Cancer Institute-Genomic Data Commons NCI-GDC (`https://portal.gdc.cancer.gov/`).

##### 1.2 Interaction networks

We used (i) STRING version 10 [1] protein-interaction network which contains high confidence functional protein-protein interactions (PPI). Self-loops and interactions with missing HGNC symbols were discarded and interaction scores were normalized (divided by 1000) to obtain a reliability score in the range $[0, 1]$. Only high confidence interactions with combined score of 0.9 or greater were selected. We also repeated our experiments using the (ii) STRING network with only experimentally verified edges, (iii) Human Protein Reference Database (HPRD) version 9 [2], as well as (iv) REACTOME version 2015 [3] PPI networks.

#### 2 Supplemental Results

##### 2.1 Distinguishing Between Different Alteration Events Contributes to the Survival Analysis

In order to demonstrate that our formulation (in this subsection we call it "multi-color") provides an advantage over a simpler approach which treats all different alteration events the same – genes are considered either altered or not, regardless of the type of alteration ("single-color"), we have computed subnetworks via both of the formulations in TCGA GBM, COAD and BRCA cohorts. We then compared the resulting plots of subnetworks' association with patient survival outcome in high and low risk groups, as explained in the methods section of the paper.

Supplementary Figure 1 shows Kaplan-Meier plots of survival probability for high and low risk groups of patients in COAD, GBM and the four BRCA subtype datasets (Basal, Luminal A, Luminal B), using STRING v10.5 interaction network. Multi-color approach achieves better separation of the curves for the two groups than the single-color approach in all datasets except Luminal B, in which both approaches detect the same subnetwork and achieve same results.

### 3 Supplementary Tables and Figures

**Supplementary Table 1.** 5 subnetworks identified by cd-CAP in multi-subnetwork mode for each cancer type: respective columns below depict the subnetwork size, depth, and the number of nodes in the subnetwork with copy number amplification (AMP), expression increase (EXP-UP) or decrease (EXP-DOWN).

| Cancer | Network# | Size | Depth | AMP | EXP-UP | EXP-DOWN |
|---|---|---|---|---|---|---|
| COAD | 1 | 6 | 206 | 1 | 5 | 0 |
| | 2 | 11 | 152 | 6 | 5 | 0 |
| | 3 | 12 | 137 | 7 | 3 | 2 |
| | 4 | 15 | 149 | 11 | 1 | 3 |
| | 5 | 15 | 223 | 2 | 10 | 3 |
| GBM | 1 | 4 | 72 | 4 | 0 | 0 |
| | 2 | 4 | 69 | 4 | 0 | 0 |
| | 3 | 9 | 67 | 9 | 0 | 0 |
| | 4 | 16 | 70 | 1 | 15 | 0 |
| | 5 | 36 | 96 | 1 | 32 | 3 |
| BRCA | 1 | 8 | 164 | 7 | 0 | 1 |
| | 2 | 10 | 332 | 1 | 9 | 0 |
| | 3 | 11 | 360 | 1 | 10 | 0 |
| | 4 | 15 | 313 | 1 | 14 | 0 |
| | 5 | 15 | 335 | 1 | 14 | 0 |

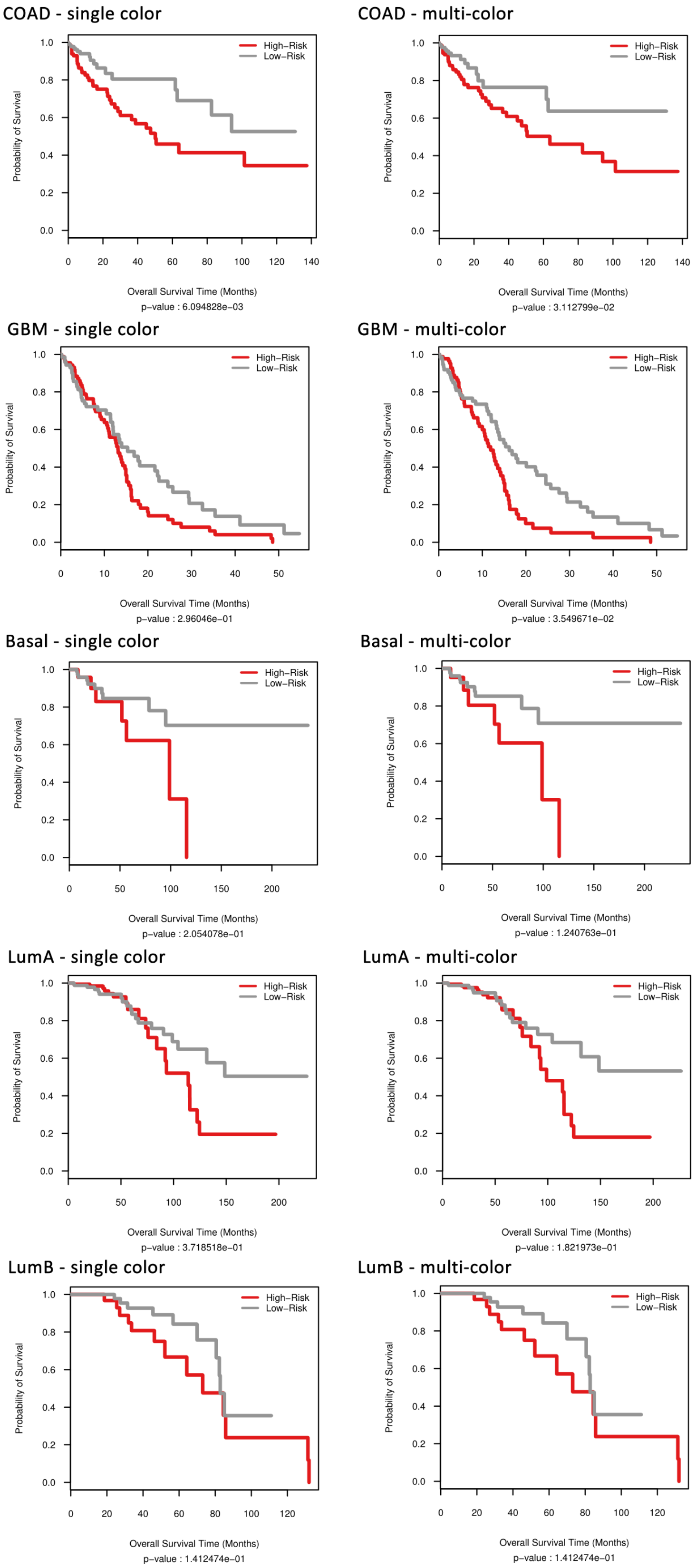


**Supplementary Figure 1.** Kaplan-Meier plot showing the association of the subnetworks with patients' clinical outcome (High Risk vs Low Risk patients), on TCGA datasets and STRING v10.5 network. The left side shows single-color formulation plots, and the right column using the multi-color formulation. The multi-color formulation achieves better separation of the curves in all datasets except LumB.

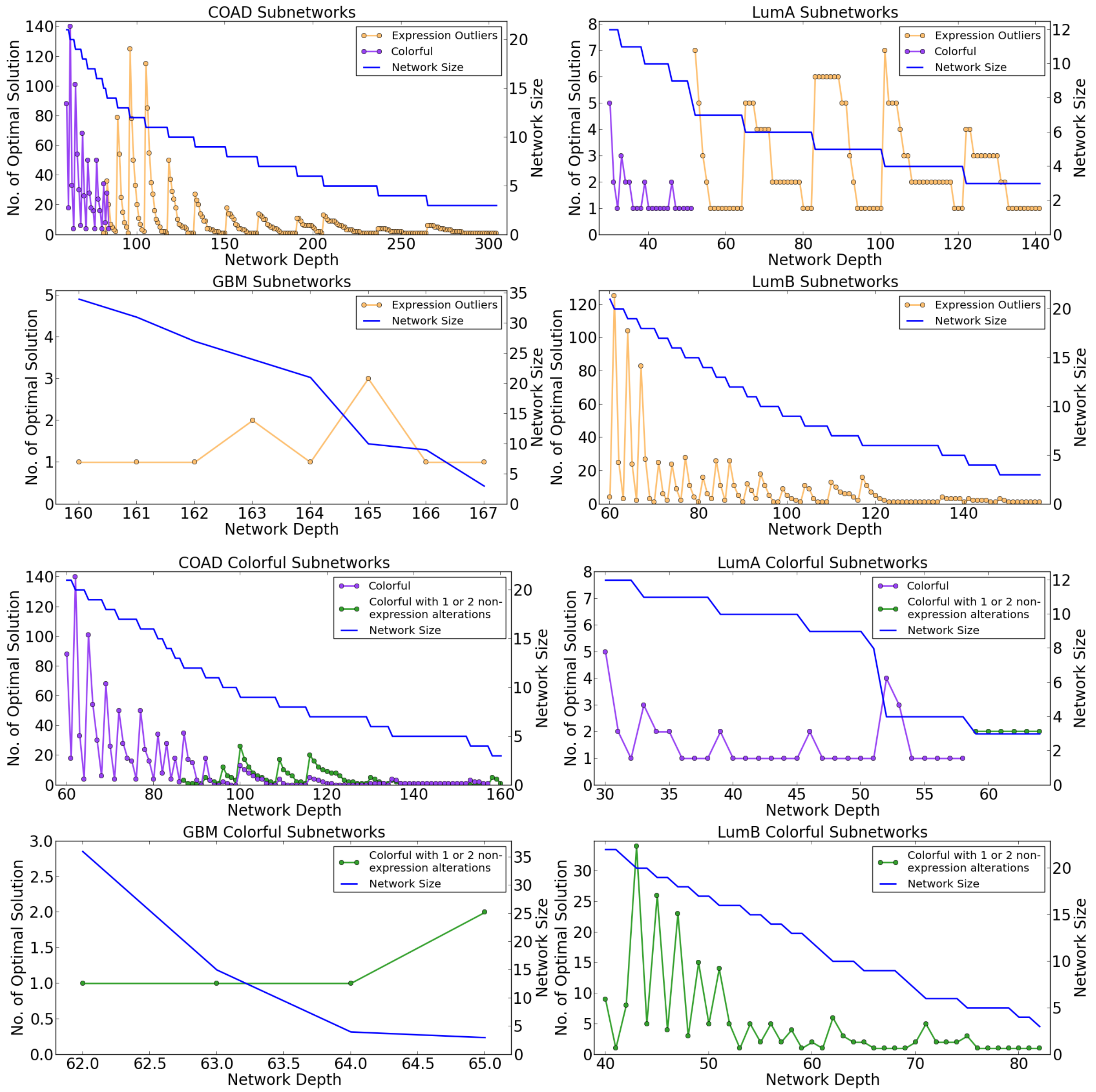


**Supplementary Figure 2.** A comprehensive view of the size of maximum subnetworks (identified through the MCSI formulation, both without and with the colorful constraint), and the number of such maximum subnetworks obtained from STRING 10 PPI network with experimentally verified edges, as a function of input network depth $t$, in each of the four TCGA datasets. The horizontal axis denotes the depth (number of patients) of the network. For the blue plot, the vertical axis (right) denotes the maximum possible network size (in terms of the number of nodes) and thus it is strictly non-increasing by definition. For the plots with different colors, the vertical axis (left) denotes the number of distinct networks with network size equal to the indicated by the blue plot. As can be seen, the red plots depict networks where all nodes have a copy number gain, the yellow plots depict networks where all nodes are expression outliers and purple plots depict colorful networks (with at least two distinct colors).

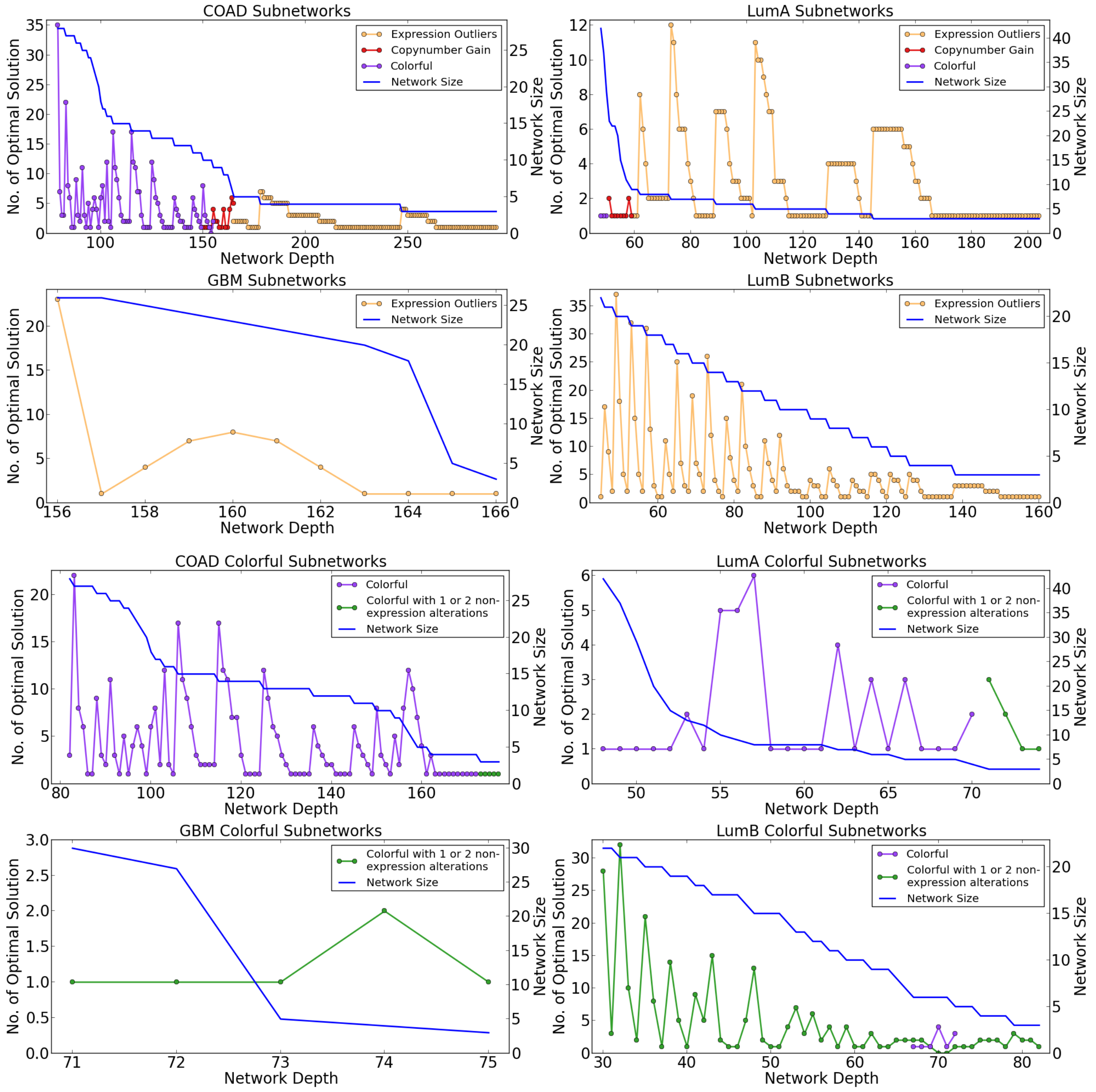


**Supplementary Figure 3.** A comprehensive view of the size of maximum subnetworks (identified through the MCSI formulation, both without and with the colorful constraint), and the number of such maximum subnetworks obtained from HPRD PPI network, as a function of input network depth $t$, in each of the four TCGA datasets. The horizontal axis denotes the depth (number of patients) of the network. For the blue plot, the vertical axis (right) denotes the maximum possible network size (in terms of the number of nodes) and thus it is strictly non-increasing by definition. For the plots with different colors, the vertical axis (left) denotes the number of distinct networks with network size equal to the indicated by the blue plot. As can be seen, the red plots depict networks where all nodes have a copy number gain, the yellow plots depict networks where all nodes are expression outliers and purple plots depict colorful networks (with at least two distinct colors).

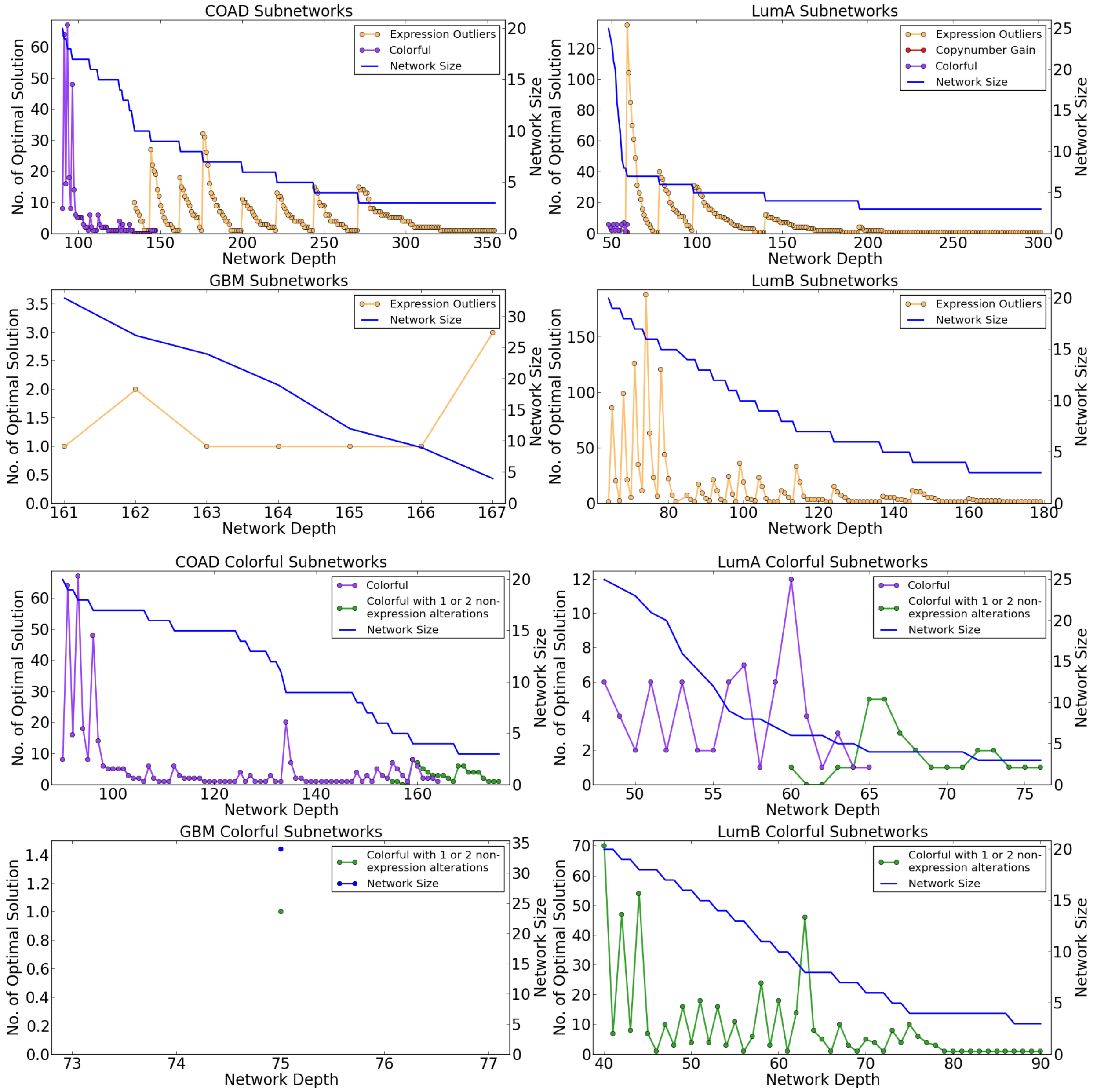


**Supplementary Figure 4.** A comprehensive view of the size of maximum subnetworks (identified through the MCSI formulation, both without and with the colorful constraint), and the number of such maximum subnetworks obtained from REACTOME PPI network, as a function of input network depth $t$, in each of the four TCGA datasets. The horizontal axis denotes the depth (number of patients) of the network. For the blue plot, the vertical axis (right) denotes the maximum possible network size (in terms of the number of nodes) and thus it is strictly non-increasing by definition. For the plots with different colors, the vertical axis (left) denotes the number of distinct networks with network size equal to the indicated by the blue plot. As can be seen, the red plots depict networks where all nodes have a copy number gain, the yellow plots depict networks where all nodes are expression outliers and purple plots depict colorful networks (with at least two distinct colors).

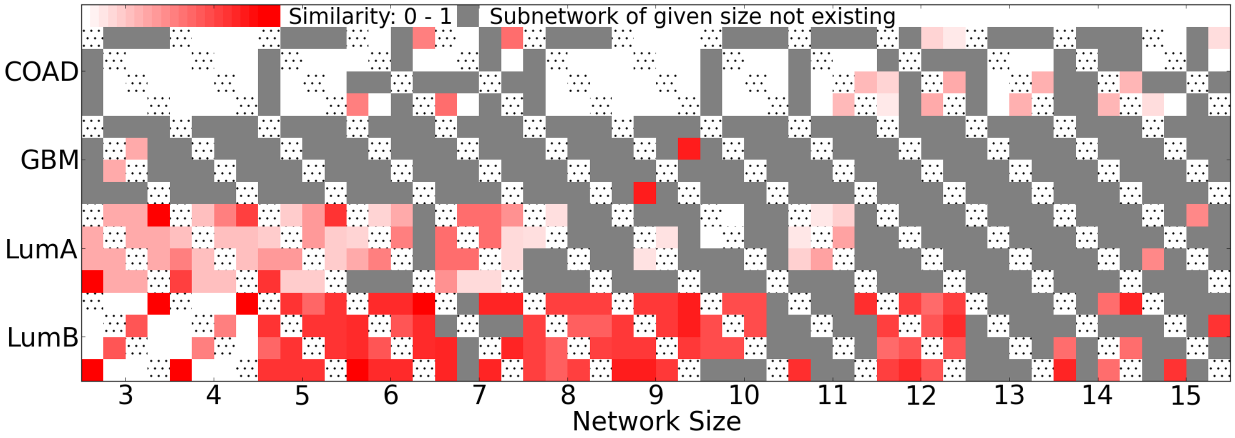


**Supplementary Figure 5.** Similarity between the candidate subnetworks (of size 3-15) identified by cd-CAP with different PPI networks as input. Each square unit corresponds to the similarity (i.e., number of overlap nodes with the same color, normalized by subnetwork size) between two maximum subnetworks of the same size (x-axis) but from different protein interaction networks (STRING, STRING with only experimentally verified edges, HPRD and REACTOME), with the exception of grey color, indicating a subnetwork of that size was not selected based on the filtering process described in Section 3.1 for one of the two protein interaction networks. Compared with the subnetworks obtained from TCGA COAD dataset, those from BRCA dataset (in particular, the Luminal B subtype) exhibit more node-overlaps, even though the node (gene) sets of the four input PPI networks are much different.

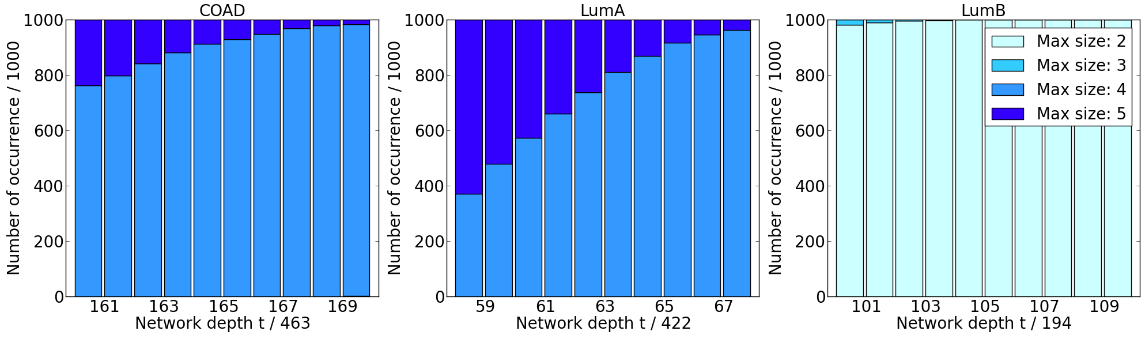


**Supplementary Figure 6.** A typical network size distribution for the maximum size subnetworks identified by cd-CAP in 1000 permutation tests with selected range of depth $t$ (notice that we do not differentiate the cases where cd-CAP return a subnetwork of size 0, 1 or 2). Compared with the subnetworks observed in real mutation profiles, those identified by cd-CAP in permutation tests (with similar or identical $t$ values) were much smaller.

**Supplementary Table 2.** Empirical p-value estimates (including the maximum and minimum largest subnetwork sizes obtained in 1000 permutation tests) for the 41 maximum-sized subnetworks identified by cd-CAP shwon in Figure 2. Compared with the subnetworks observed in real mutation profiles, those identified by cd-CAP in permutation tests (with identical $t$ values) were much smaller, implying a p-value of $< 0.001$ for each of the colored subnetworks presented in Figure 2.

| Dataset | Depth | Size | Max Size Simulated | Minmax Size Simulated | p-value |
|---|---|---|---|---|---|
| COAD | $t = 283$ | 4 | $\leq 2$ | $\leq 2$ | $< 0.001$ |
| COAD | $t = 234$ | 6 | 3 | $\leq 2$ | $< 0.001$ |
| COAD | $t = 224$ | 7 | 3 | $\leq 2$ | $< 0.001$ |
| COAD | $t = 163$ | 9 | 5 | 4 | $< 0.001$ |
| COAD | $t = 161$ | 12 | 5 | 4 | $< 0.001$ |
| COAD | $t = 159$ | 15 | 5 | 4 | $< 0.001$ |
| COAD | $t = 156$ | 18 | 5 | 4 | $< 0.001$ |
| COAD | $t = 153$ | 22 | 6 | 4 | $< 0.001$ |
| COAD | $t = 149$ | 27 | 6 | 4 | $< 0.001$ |
| COAD | $t = 147$ | 30 | 6 | 4 | $< 0.001$ |
| GBM | $t = 167$ | 11 | $\leq 2$ | $\leq 2$ | $< 0.001$ |
| GBM | $t = 165$ | 20 | $\leq 2$ | $\leq 2$ | $< 0.001$ |
| GBM | $t = 164$ | 27 | $\leq 2$ | $\leq 2$ | $< 0.001$ |
| GBM | $t = 163$ | 30 | $\leq 2$ | $\leq 2$ | $< 0.001$ |
| LumA | $t = 301$ | 3 | $\leq 2$ | $\leq 2$ | $< 0.001$ |
| LumA | $t = 259$ | 4 | $\leq 2$ | $\leq 2$ | $< 0.001$ |
| LumA | $t = 178$ | 5 | $\leq 2$ | $\leq 2$ | $< 0.001$ |
| LumA | $t = 120$ | 6 | 4 | 3 | $< 0.001$ |
| LumA | $t = 103$ | 7 | 4 | 3 | $< 0.001$ |
| LumA | $t = 90$ | 8 | 4 | 3 | $< 0.001$ |
| LumA | $t = 60$ | 10 | 5 | 4 | $< 0.001$ |
| LumA | $t = 57$ | 11 | 5 | 4 | $< 0.001$ |
| LumA | $t = 56$ | 15 | 5 | 4 | $< 0.001$ |
| LumA | $t = 51$ | 28 | 6 | 4 | $< 0.001$ |
| LumA | $t = 50$ | 33 | 6 | 4 | $< 0.001$ |
| LumA | $t = 49$ | 40 | 6 | 4 | $< 0.001$ |
| LumA | $t = 48$ | 46 | 6 | 4 | $< 0.001$ |
| LumB | $t = 170$ | 3 | $\leq 2$ | $\leq 2$ | $< 0.001$ |
| LumB | $t = 159$ | 4 | $\leq 2$ | $\leq 2$ | $< 0.001$ |
| LumB | $t = 143$ | 5 | $\leq 2$ | $\leq 2$ | $< 0.001$ |
| LumB | $t = 136$ | 6 | $\leq 2$ | $\leq 2$ | $< 0.001$ |
| LumB | $t = 131$ | 7 | $\leq 2$ | $\leq 2$ | $< 0.001$ |
| LumB | $t = 124$ | 8 | $\leq 2$ | $\leq 2$ | $< 0.001$ |
| LumB | $t = 115$ | 9 | $\leq 2$ | $\leq 2$ | $< 0.001$ |
| LumB | $t = 109$ | 10 | $\leq 2$ | $\leq 2$ | $< 0.001$ |
| LumB | $t = 104$ | 11 | 3 | $\leq 2$ | $< 0.001$ |
| LumB | $t = 99$ | 12 | 3 | $\leq 2$ | $< 0.001$ |
| LumB | $t = 94$ | 13 | 3 | $\leq 2$ | $< 0.001$ |
| LumB | $t = 90$ | 14 | 3 | $\leq 2$ | $< 0.001$ |
| LumB | $t = 83$ | 16 | 3 | $\leq 2$ | $< 0.001$ |
| LumB | $t = 80$ | 17 | 3 | $\leq 2$ | $< 0.001$ |
| LumB | $t = 77$ | 18 | 3 | $\leq 2$ | $< 0.001$ |
| LumB | $t = 70$ | 20 | 4 | 3 | $< 0.001$ |

**Supplementary Table 3.** Empirical p-value estimates (including the maximum and minimum largest subnetwork sizes obtained in 1000 permutation tests) for the 28 maximum-sized subnetworks identified by cd-CAP in the STRING v10 (high confidence edges only, out of 104 in all PPI networks) PPI network, shwon in Figure 3A-D. Compared with the subnetworks observed in real mutation profiles, those identified by cd-CAP in permutation tests (with identical $t$ values) were much smaller, implying a p-value of $< 0.001$ for each of the colored subnetworks presented in Figure 3A-D.

| Dataset | Depth | Size | Max Size Simulated | Minmax Size Simulated | p-value |
|---|---|---|---|---|---|
| COAD | $t = 177$ | 3 | | | $< 0.001$ |
| COAD | $t = 172$ | 3 | | | $< 0.001$ |
| COAD | $t = 171$ | 5 | | | $< 0.001$ |
| COAD | $t = 167$ | 6 | | | $< 0.001$ |
| COAD | $t = 160$ | 10 | $\leq 2$ | $\leq 2$ | $< 0.001$ |
| COAD | $t = 158$ | 13 | | | $< 0.001$ |
| COAD | $t = 156$ | 16 | | | $< 0.001$ |
| COAD | $t = 153$ | 19 | | | $< 0.001$ |
| COAD | $t = 146$ | 28 | | | $< 0.001$ |
| GBM | $t = 75$ | 9 | $\leq 2$ | $\leq 2$ | $< 0.001$ |
| GBM | $t = 74$ | 53 | | | $< 0.001$ |
| LumA | $t = 79$ | 3 | | | $< 0.001$ |
| LumA | $t = 74$ | 4 | | | $< 0.001$ |
| LumA | $t = 68$ | 5 | | | $< 0.001$ |
| LumA | $t = 62$ | 6 | $\leq 2$ | $\leq 2$ | $< 0.001$ |
| LumA | $t = 60$ | 7 | | | $< 0.001$ |
| LumA | $t = 55$ | 12 | | | $< 0.001$ |
| LumA | $t = 54$ | 16 | | | $< 0.001$ |
| LumB | $t = 91$ | 3 | | | $< 0.001$ |
| LumB | $t = 87$ | 4 | | | $< 0.001$ |
| LumB | $t = 85$ | 5 | | | $< 0.001$ |
| LumB | $t = 76$ | 8 | | | $< 0.001$ |
| LumB | $t = 74$ | 9 | | | $< 0.001$ |
| LumB | $t = 72$ | 10 | $\leq 2$ | $\leq 2$ | $< 0.001$ |
| LumB | $t = 68$ | 12 | | | $< 0.001$ |
| LumB | $t = 65$ | 14 | | | $< 0.001$ |
| LumB | $t = 62$ | 15 | | | $< 0.001$ |
| LumB | $t = 60$ | 16 | | | $< 0.001$ |
